# Long-term experimental evolution reveals purifying selection on piRNA-mediated control of transposable element expression

**DOI:** 10.1101/666693

**Authors:** Ulfar Bergthorsson, Caroline J. Sheeba, Anke Konrad, Tony Belicard, Toni Beltran, Vaishali Katju, Peter Sarkies

## Abstract

Transposable elements (TEs) are an almost universal constituent of eukaryotic genomes. In animals, Piwi-interacting small RNAs (piRNAs) and repressive chromatin often play crucial roles in preventing TE transcription and thus restricting TE activity. Nevertheless, TE content varies widely across eukaryotes and the dynamics of TE activity and TE silencing across evolutionary time is poorly understood. Here we used experimentally evolved populations of *C. elegans* to study the dynamics of TE expression over 400 generations. The experimental populations were evolved at three different population sizes to manipulate the efficiency of natural selection versus genetic drift. We demonstrate increased TE expression relative to the ancestral population, with the largest increases occurring in the smallest populations. We show that the transcriptional activation of TEs within active regions of the genome is associated with failure of piRNA-mediated silencing, whilst desilenced TEs in repressed chromatin domains retain small RNAs. Additionally, we find that the sequence context of the surrounding region influences the propensity of TEs to lose silencing through failure of small RNA-mediated silencing. Together, our results show that natural selection in *C. elegans* is responsible for maintaining low levels of TE expression, and provide new insights into the epigenomic features responsible.

## Introduction

Transposable elements (TEs) are almost ubiquitous across eukaryotic genomes (Chuong et al., 2017). Their ability to replicate independently of the host genome, coupled with the existence of multiple copies liable to ectopic recombination means they present a potential threat to genome stability. Moreover, TEs pose a threat to genome function as new integrations can disrupt genes or gene regulatory elements. As a result, organisms have evolved sophisticated control strategies, which protect the genome from TE proliferation. Across eukaryotes, short (20-33 nucleotides) small RNAs play an important role in the suppression of TE activity. Within animals, Piwi-interacting small RNAs (piRNAs) are paramount in the TE defence armoury (Siomi et al., 2011). piRNAs are produced from defined genomic loci named piRNA clusters and after processing, associate with the Piwi subfamily of argonaute proteins (Brennecke et al., 2007). They recognise TEs through sense-antisense base pairing and target TEs for transcriptional and post-transcriptional silencing(Siomi et al., 2011). In many model organisms, piRNAs are essential for fertility through their role in controlling TE proliferation in the germline (Weick and Miska, 2014)

The nematode *Caenorhabditis elegans* is a well-established model for small-RNA mediated silencing. piRNAs in *C. elegans* are unusual in that the two piRNA clusters on Chromosome IV are composed of individual RNA polymerase II (RNA pol II) transcription loci where each piRNA has its own upstream motif (Batista et al., 2008; Das et al., 2008; Ruby et al., 2006; Wang and Reinke, 2008). piRNA clusters are located within H3K27me3-rich chromatin, which, together with cis-acting RNA pol II pausing sequences downstream of the piRNA, enforce production of ∼28 nucleotide piRNA precursors(Beltran et al., 2019). piRNA precursors are further trimmed to result in mature 21 nucleotide piRNAs with a Uracil as the first nucleotide (21U-RNAs), which associate with the *C. elegans* Piwi protein PRG-1 (Batista et al., 2008; Das et al., 2008; Wang and Reinke, 2008). Downstream of PRG-1, piRNA silencing relies on a nematode-specific class of secondary small RNAs known as 22G-RNAs(Das et al., 2008). 22G-RNA synthesis is carried out by RNA-dependent RNA polymerases using the target RNA as a template, following initiation by piRNA target recognition(Pak and Fire, 2007). 22G-RNAs bind to Argonaute proteins and lead to transcriptional and post-transcriptional silencing of target RNAs (Yigit et al., 2006). Additionally 22G-RNAs can be transmitted transgenerationally (Buckley et al., 2012) and as a result piRNA-initiated silencing can persist for many generations even after piRNAs themselves are removed by mutating PRG-1 (Ashe et al., 2012; Luteijn et al., 2012; Shirayama et al., 2012). Consequently whilst removal of piRNAs alone has mild effects on TE expression, combining mutations of PRG-1 with mutations disrupting the 22G-RNA biogenesis machinery leads to reactivation of several TEs (de Albuquerque et al., 2015; Phillips et al., 2015)

Despite the universality of TEs across eukaryotes, there is striking variability both in TE content and TE expression across species. Several interacting factors have been proposed to account for this. First, silencing mechanisms differ between organisms. For example, the entire piRNA pathway has been lost independently multiple times in nematodes (Sarkies et al., 2015), and was lost in parasitic flatworms (Fontenla et al., 2017; Skinner et al., 2014) and in dust mites (Mondal et al., 2018). Second, it is possible that some TEs may have beneficial consequences through their ability to act as reservoirs for evolutionary novelty. For example up to 60% of human-specific enhancers may be TE-derived (Rebollo et al., 2011) and TE insertions have been proposed to substantially rewire the human immune cell transcriptome (Imbeault et al., 2017). TEs themselves may also be co-opted into developmental programs. For example, transcription of L1 RNA is observed at the 2-cell stage in mouse embryogenesis where it may have a direct role in coordinating gene expression programs (Percharde et al., 2018). Furthermore, TEs may serve as a genome-wide source of regulatory elements (Chuong et al., 2017). Despite these examples, TEs are overall considered to be detrimental to fitness and beneficial TE insertions appear overrepresented due to the effects of natural selection in weeding out deleterious insertions (Simonti et al., 2017). The diversity of TEs across evolution may thus reflect population genetics factors such as population structure and effective population size. For example, even moderately deleterious TE insertions might become fixed in very small populations as the intensity of selection decreases with reduced population size. In agreement with this model, a recent large-scale study across nematodes concluded that genetic drift was likely responsible for differences in TE content across nematodes (Szitenberg et al., 2016).

In the context of these potential models to explain the diversity in TE content, it is important to understand the extent to which the balance between TE expression and TE regulation is under selection. One way to study this is to use a mutation accumulation (MA) framework in which replicate lines descended from a single common ancestor are propagated under a regime of drastic population bottlenecks for several hundred generations (Halligan and Keightley, 2009; Katju and Bergthorsson, 2019). The maintenance of these lines at a minimal population size attenuates the efficacy of selection, thereby enabling the accumulation of a large, unbiased sample of spontaneous mutations under conditions of genetic drift which can subsequently be identified and their fitness effects investigated. Previously, MA lines have been used to investigate the rate and spectrum of TE copy-number changes in *Saccharomyces cerevisiae*, *C. elegans* and *Drosophila melanogaster*, thus providing estimates of the rate of TE transposition (Adrion et al., 2017; Bast et al., 2019; Bégin and Schoen, 2006, 2007). However, TE expression is not necessarily directly linked to TE copy-number and may have independent fitness consequences. The opportunity to interrogate genome-wide RNA expression has been exploited to investigate the effect of mutations on protein-coding gene expression divergence (Denver et al., 2005; Landry et al., 2007; Rifkin et al., 2005). Here, we extend this approach to investigate the effect of spontaneous mutations on TE expression divergence.

We created spontaneous MA lines of *C. elegans* that were descended from a single worm ancestor and propagated for ∼400 generations under three population size treatments of *N* = 1, 10 and 100 individuals per generation (Katju et al., 2015). The varying population size treatment in the experiment permitted a manipulation of the strength of selection, with the *N* =1 lines evolving under close to neutral conditions (minimal selection) and an incremental increase in the strength of selection with increasing population size. We employed this framework to investigate how TE expression evolves under conditions of near neutrality and under the influence of increasing selection intensity. We show that overall TE expression increases in MA lines with the smallest population size. We further show that expression increase results in part from failure of piRNA-mediated silencing. Intriguingly, differences in the responses of different TEs to reduced piRNA-mediated silencing depend on the chromatin environment of the TE loci, such that TEs in repressed chromatin domains largely remain silent due to epigenetic memory imparted by 22G-RNAs, whilst in active chromatin domains, increased TE expression is much more likely to occur. Together our results demonstrate for the first time that robust control of TE expression is under selection in animals. Importantly further, our results provide new insight into how the chromatin environment interacts with piRNA-mediated silencing to control TE expression.

## Results

### Relaxed selection leads to increased TE expression

In order to assess the effect of selection on TE expression, we generated mutation accumulation (MA) lines and propagated them by randomly selecting *N* individuals at each generation, where *N* was either 1, 10 or 100 (Katju et al., 2015). Henceforth we refer to the three conditions as *N*.1, *N*.10 or *N*.100 (Figure 1A). After 409 generations, we isolated RNA and performed RNA sequencing to investigate TE expression. The fundamental difference between these conditions is the size of the bottleneck that the population is subjected to in each generation. However, it is inevitable that the population density will also vary between the three population size treatments, resulting in environmental differences that could introduce variation in gene expression among the experimental lines maintained at these three population sizes. To counteract this, all lines were maintained in similar conditions including no population bottleneck differences for three generations prior to RNA extraction. Therefore, the most likely explanation for any differences in TE expression that we observe is the different strength of purifying selection between the three population size treatments during MA.

**Figure 1.**
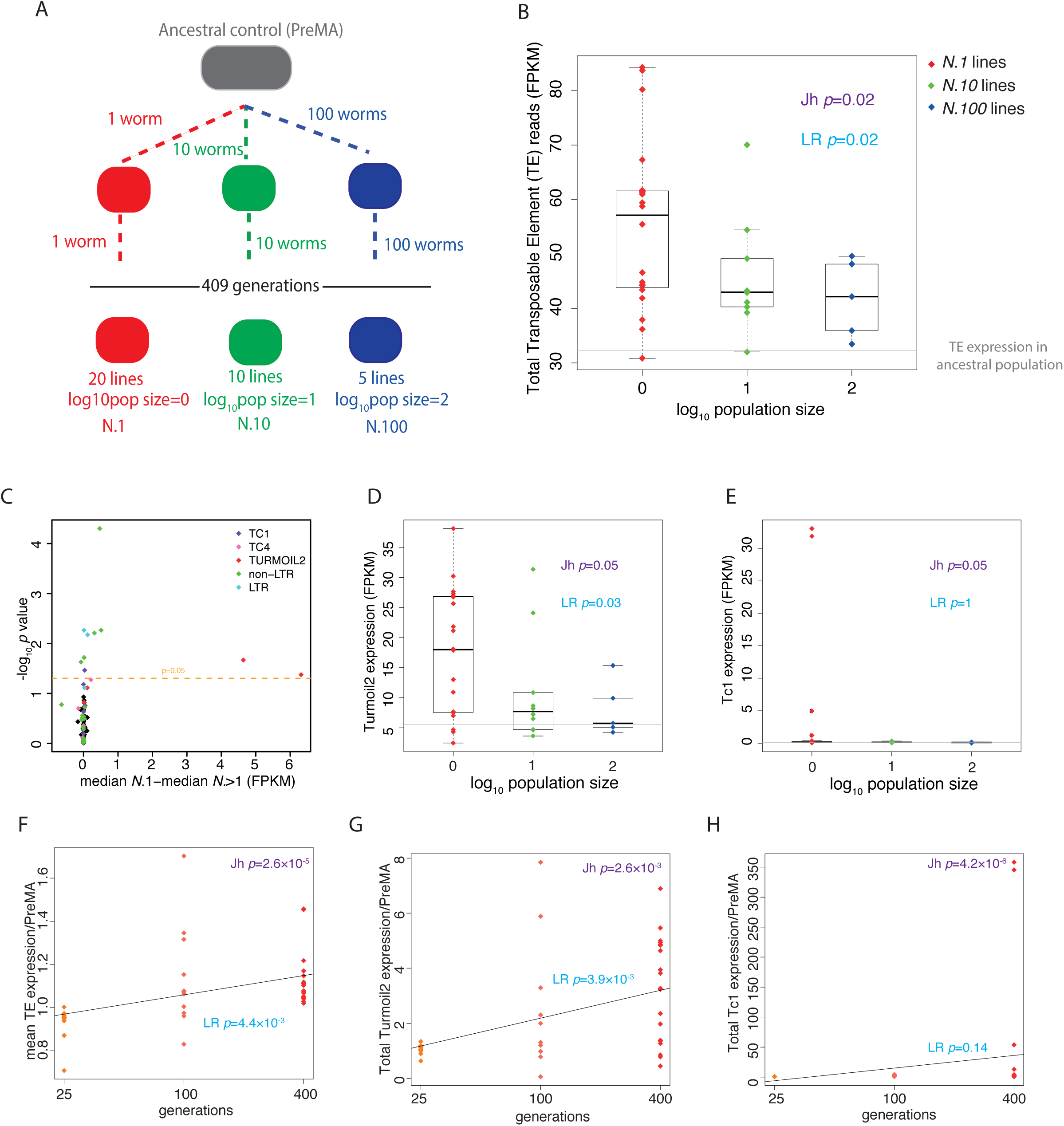
Increased expression of transposable elements in mutation accumulation lines. **A** Diagram of the mutation accumulation (MA) experimental design with different population sizes. **B** Overall transposable element expression in MA lines as a function of population size. **C** Volcano plot for individual TE genes coloured according to family. The x axis shows the median difference between *N*.1 and combined *N*.10 and *N*.100 lines such that higher values correspond to higher expression in smaller population sizes. The p-value is a Wilcoxon unpaired test comparing the median in *N*.1 lines to *N*.10 and *N*.100. **D** Expression changes in Turmoil2 elements. The box shows interquartile range with a line at median and the whiskers extend to the furthest point ≤ 1.5 times the interquartile range from the median. **E** Expression changes in Tc1 elements. Boxplot as in D. **F** Total TE expression after different numbers of generations of mutation accumulation in lines with a population size of 1. **G** Turmoil2 expression after different numbers of generations of mutation accumulation with a population size of 1. **H** Tc1 expression after different numbers of generations of mutation accumulation with a population size of 1.

MA lines from all three population size treatments showed an increase in total TE expression relative to the pre-MA ancestral control; moreover, *N*.1 had higher total TE expression than *N*.10 or *N*.100. Across increasing population sizes, we observed a monotonic decrease in total TE expression (Jonckheere test for ordered medians; henceforth Jh, *p* = 0.02; Figure 1B). Similarly, linear regression analysis showed a significant negative correlation between increasing population size and TE expression (linear regression; henceforth LR, *p* = 0.02 Figure 1B). The mean expression across all changes in TEs normalized to the pre-MA ancestral control showed a significant tendency to decrease as the population size increased (Jh, *p* = 0.003; LR, *p* = 0.02; Supplemental Figure 1A).

Protein-coding gene expression diverges during MA in a variety of model organisms including *C. elegans* (Denver et al., 2005; Rifkin et al., 2005; Landry et al., 2007; Hodgins-Davis et al., 2015). In addition to an overall increase, TE expression also appears to show broader distributions in the *N*.1 lines compared to the *N*.10 and *N*.100 lines (Supplemental Figure 1B). To test this directly, we estimated the variation in the expression of each individual TE and each individual gene. To control for potential changes in the mean expression, which can affect noise, we calculated the Fano factor [var(x)/mean(x)] within *N*.1, *N*.10 and *N*.100 lines separately. Fano factors for TEs and genes were higher in *N*.1 lines than in *N*.10 or *N*.100 lines (TEs: Wilcoxon paired test *p* = 0.015 and 0.004 for *N*.10 and *N*.100; genes: Wilcoxon paired test *p* < 1×10^-16^ for both *N*.10 and *N*.100; Supplemental Figures 1C and D). To control for the possibility that the larger number of *N*.1 lines might lead to higher variance, we calculated TE Fano factors from 1,000 subsets of five *N*.1 lines and all 252 subsets of five *N*.10 lines and compared these to the five *N*.100 lines. This showed the same trend as the full dataset (Supplemental Figure 1E). To further investigate the variation in expression, we calculated the total variance in the change in expression of all TEs or all genes between each line and the pre-MA ancestral control. Variance in the differences in both TE and gene expression increased with smaller population sizes (Jh, *p* = 0.008 and 0.009, respectively) (Supplemental Figure 1 F,G). However, importantly, there was no correlation in overall variance between TEs and genes in the same line (Supplemental Figure 1H), showing that TE expression and gene expression diverge independently.

In order to understand loss of repression of TE expression in more detail, we classified TEs into different families using RepeatMasker annotations and compared the median TE expression for each TE family across different lines. TE expression was compared between *N*.1 and *N*>1 lines and tested for monotonic median increase with decreasing population size. As expected, the majority of TEs that showed a significant difference between *N*.1 and combined *N*.10 and *N*.100 lines had increased expression in *N*.1 compared to the other lines (Figure 1C and Supplemental Figure 2). However, individual TEs displayed different patterns of expression change. Some TEs, notably the DNA transposon Turmoil2, showed more consistent increases across the *N*.1 lines compared to the *N*.10 and *N*.100 lines (Figure 1D). Indeed, the majority of the total effect on TE expression seen in Figure 1B could be attributed to one TE family, the Turmoil2 TEs, which showed a large expression increase across the majority of the *N*.1 lines (Figure 1C, D). Contrastingly some TEs, notably the Mariner family DNA transposon Tc1, showed a burst-like pattern of expression where a large increase in expression was observed in a few *N*.1 lines while retaining low expression in the remainder *N*.1 lines as well as the *N*>1 lines (Figure 1E).

We next investigated the timecourse of TE desilencing during propagation of the MA lines. We performed gene expression analysis by RNA-Seq on 11 *N*.1 lines at 25 and 100 MA generations. Median total TE expression showed a highly significant increase with increasing numbers of generations (Jh, *p* = 2.6×10^-5^; Figure 1F). Linear regression analysis confirmed a positive relationship between increased numbers of generations and increased TE expression (LR, *p* = 4.4×10^-3^; Figure 1F). Different TEs showed different kinetics of desilencing. Turmoil2 showed a positive relationship between the number of MA generations and expression (LR *p* = 3.9×10^-3^) and a monotonic increase in median expression (Jh, *p* = 2.6×10^-3^; Figure 1G). Contrastingly, Tc1 desilencing did not show a positive linear relationship between the number of MA generations and expression (LR, *p* = 0.14; Figure 1H) though there was a significant increase in median expression (Jh, *p* = 4.2×10^-6^).

We further investigated whether the expression of TEs in individual MA lines correlated with the expression of other TEs. The majority of TEs showed little correlation with the expression of other TEs (Figure 2A, B). The TEs with the most statistically significant increases in expression, Turmoil2 and non-LTR retrotransposons of the LINE2 family (Figure 1C and Supplemental Figure 2) clustered together (Figure 2B) suggesting coregulation of these TE families despite their different mechanisms of replication.

**Figure 2.**
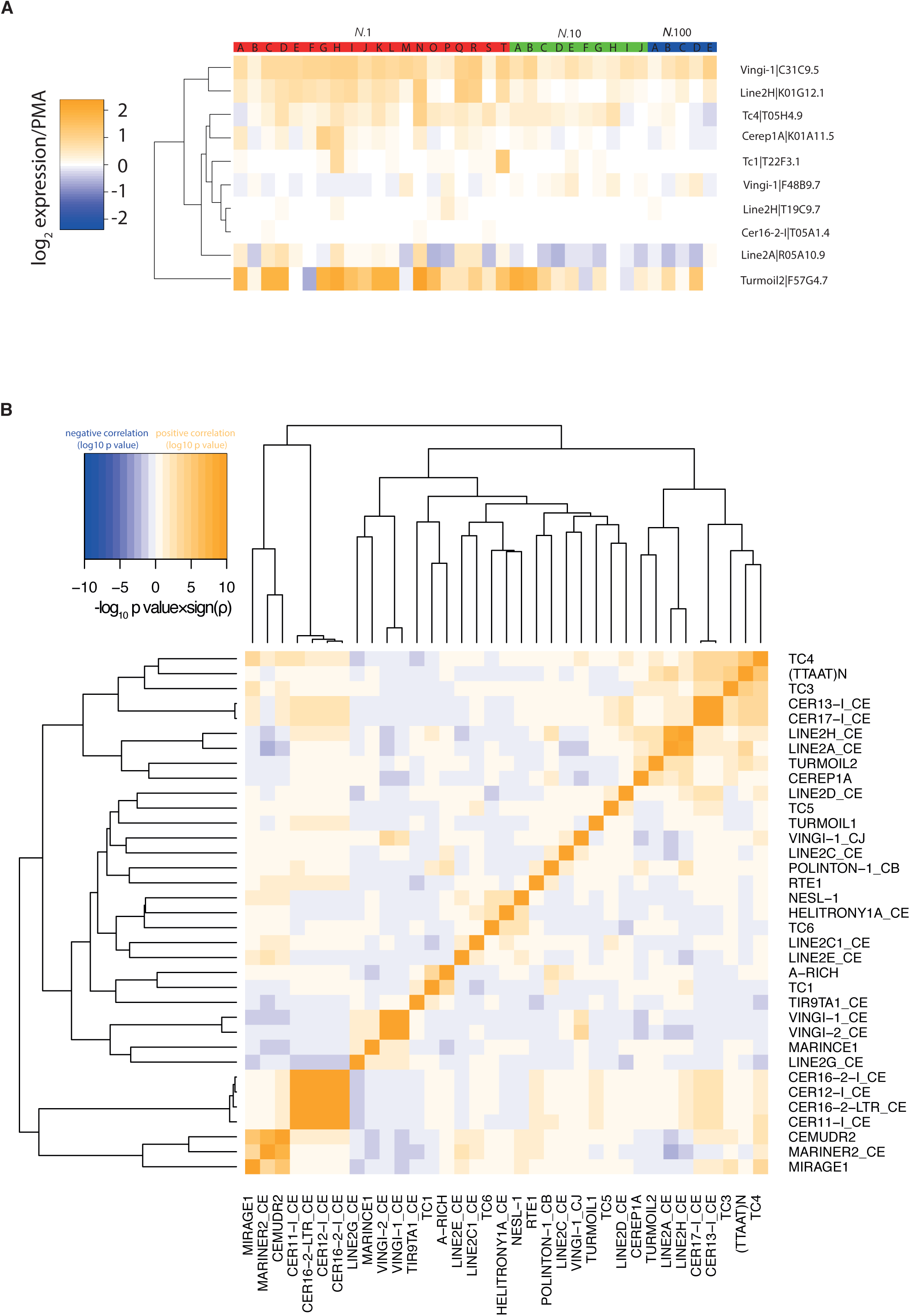
A heatmap of expression of individual TE elements illustrating significant differences in expression between different population size treatments. **B** A heatmap illustrating the significance in correlation in expression between different TEs across all the MA lines. The colour intensity shows the significance of the Spearman’s rank correlation coefficient with blue showing a negative correlation and orange a positive correlation.

### Expression of TEs is weakly associated with increased copy-number

TEs are capable of replicating independently of the host genome and thus their copy-number might change across MA lines. We sequenced the genomes of the MA lines after 400 generations and mapped the reads to consensus TE sequences thereby obtaining estimates of copy-number variation (CNV) for each TE family. Median TE copy-number increased with decreasing population size (Jh, *p* = 8.0×10^-4^; LR, *p* = 4.2×10^-3^ Figure 3A). Across all lines, there was a significant positive correlation (LR, *p* = 0.01) between increased copy-number and increased expression, although this was driven by *N.*1 lines and not apparent in the *N.*10 and *N.*100 lines (Figure 3B). Furthermore, Turmoil2 elements, which exhbitied the largest changes in expression, displayed no correlation between expression changes and copy-number increases (Figure 3C). We conclude that increased TE copy-number was not the primary cause of increased TE expression. Moreover, increased expression of specific TEs did not always lead to increased copy-number.

**Figure 3.**
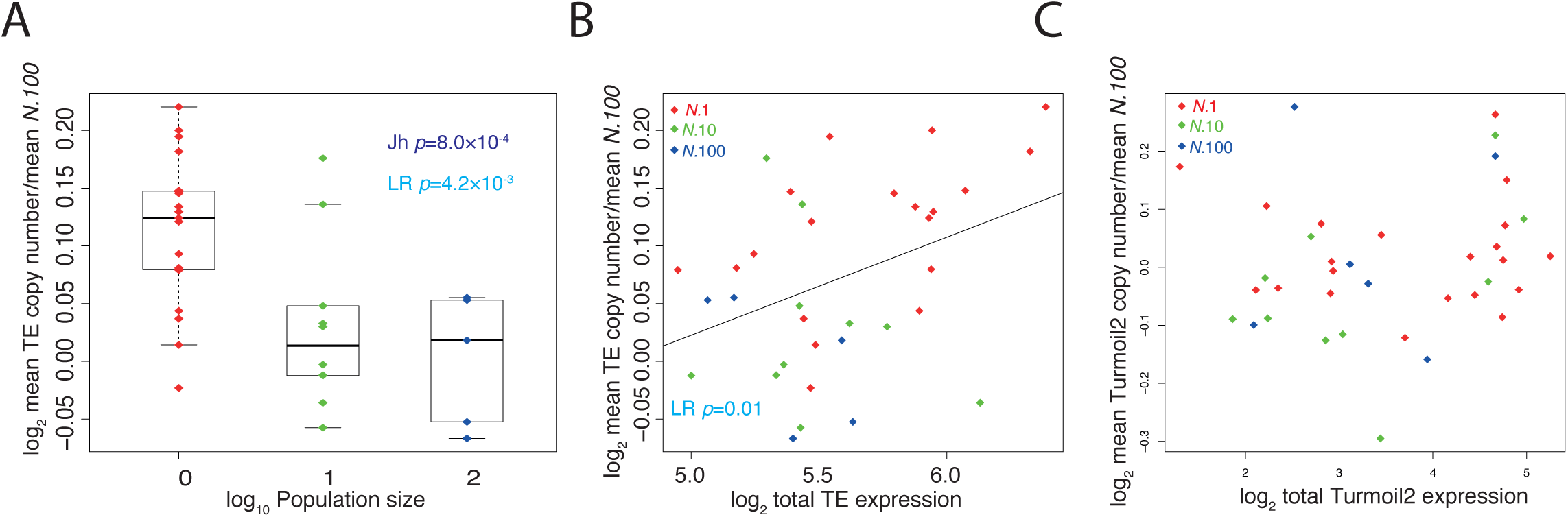
Relationship between TE copy-number and expression in MA lines. **A** Increase in TE copy-number is associated with stronger genetic drift (weaker selection). Data shows mean of TE copy-number across all TEs normalized to the mean across all lines. **B** Correlation between total change in TE copy-number across all TEs and total TE expression. **C** Lack of correlation between Turmoil2 RNA levels and copy-number changes in MA lines.

### Alterations in small RNA levels are associated with TE expression changes

We investigated whether changes in regulation of TE expression could explain the loss of silencing observed during mutation accumulation. In *C. elegans*, piRNAs and 22G-RNAs are important small RNA classes involved in TE silencing (de Albuquerque et al., 2015; Bagijn et al., 2012; Das et al., 2008; Phillips et al., 2015). To test whether piRNAs are important in the loss of silencing of TEs in the *N*.1 lines, we remapped recently published cross-linking immunoprecipitation (CLIP) data (Shen et al., 2018) to identify TE transcripts that are bound by piRNA-Piwi complexes *in vivo*. Approximately 25% of TE transcripts with RNA-Seq reads were targeted by piRNAs. TEs targeted by piRNAs showed a statistically significant increase in total expression in the *N*.1 lines (Jh, *p* = 0.02; LR, *p* = 0.03; Figure 4A) whilst TEs that were not targeted by piRNAs were not significantly altered (Jh, *p* = 0.47; LR, *p* = 0.27; Figure 4B).

**Figure 4.**
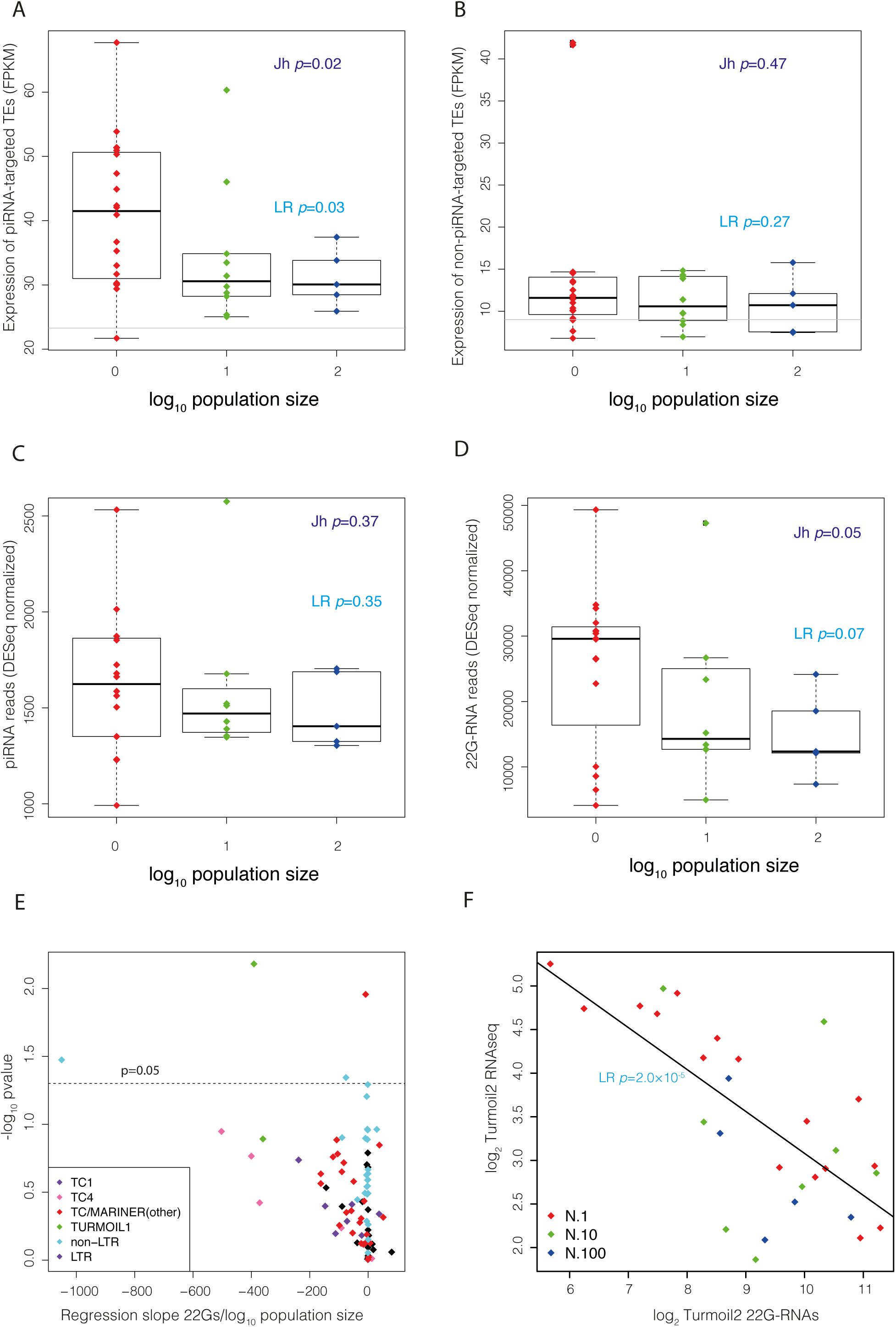
Perturbed 22G-RNAs are associated with changes in TE expression in MA lines. **A and B** Comparison of differences in TE expression in MA lines of different population sizes for piRNA targeted and non-piRNA targeted TEs, respectively. **C** Total piRNA levels in MA lines across different population sizes. **D** Levels of 22G-RNAs mapping to all TEs in MA lines at different population sizes. **E** Volcano plot showing the gradient of change in 22G levels for different TEs relative to the population size against their *p*-values on the y-axis. The gradient of change was estimated by linear regression. Negative values on the x-axis correspond to higher 22G-RNA levels in lines with smaller population sizes. **F** 22G-RNAs mapping to Turmoil2 plotted against Turmoil2 expression across all MA lines.

We tested whether defective piRNA-mediated silencing might account for increased expression of TEs targeted by piRNAs. piRNAs in *C. elegans* are expressed from individual promoters as a result of RNA polymerase II transcription (Billi et al., 2013; Cecere et al., 2012; Gu et al., 2012). We considered two potential mechanisms that might give rise to altered piRNAs. One possibility is that mutations in the piRNA sequences might occur in individual lines, which might affect their ability to recognise transposable elements and thus interfere with TE silencing. There was a trend for *N*.1 lines to have more mutations in piRNAs than *N*.10 or *N*.100 lines (Supplemental Figure 3). However, the trend was not significant (Jh, *p* = 0.12). Moreover, on average, we identified only 1.1 piRNA sequences with mutations across the *N*.1 lines. Thus, we conclude that the increase in TE expression is unlikely to be related to mutations in specific piRNAs.

We next considered the expression of individual piRNA loci. We identified piRNA loci with significantly altered median expression between *N*.1 lines and *N*.10 and *N*.100 lines combined using the Wilcoxon unpaired test. A small percentage of piRNA loci showed significant differences but overall there was no trend for these loci to show reduced expression in the *N*.1 lines. Indeed, these loci were more likely to have higher expression in the *N*.1 lines (Figure 4C). Thus, changes in piRNA expression are unlikely to explain the changes we observed in TE expression.

22G-RNAs act downstream of piRNAs to bring about target silencing(Das et al., 2008). Surprisingly, although 22G-RNAs silence TEs, we found that the total levels of 22G-RNAs mapping to TEs were increased in lines with smaller population sizes, although this increase was on the border of significance (Jh, *p* = 0.05; LR, *p* = 0.07; Figure 4D). To examine this in more detail, we analysed 22G-RNAs at individual TEs. The majority of TEs showing significantly increased 22G-RNAs in *N*.1 lines were non-LTR transposons (Figure 4E). Notably, these TEs were not significantly elevated in transcript levels in the *N*.1 lines (Figure 1C; Supplemental Figure 2). Thus, the global increase in 22G-RNA levels does not relate directly to the increased transcript levels of TEs in *N*.1 lines.

To compare directly how increased expression of TEs relates to 22G-RNA mediated silencing, for each TE we compared 22G-RNA levels in lines with increased expression to 22G-RNA levels in lines without increased expression. The only TE showing alterations in both expression and 22G-RNA levels was Turmoil2. The expression changes of Turmoil2 relative to the ancestral control across all lines correlated inversely with changes in 22G-RNA levels (LR, *p* = 2.0×10^-5^, Figure 4F). Thus, the increased transcript levels of Turmoil2 elements is associated with reduced 22G-RNAs. We conclude that the increased transcript levels of some TEs may be caused by reduced small RNA-mediated silencing, but that some increases in TE transcript levels occur independently of 22G-RNA changes.

### Chromatin environment is associated with the relationship between 22G-RNA levels and TE expression

22G-RNAs interact with chromatin modifying factors to control expression of TEs (McMurchy et al., 2017). piRNA-mediated silencing has been directly linked to the generation of H3K9me2/3 marked nucleosomes (“classical heterochromatin”) and this has been proposed to be important for transcriptional silencing induced by piRNAs (Ashe et al., 2012; McMurchy et al., 2017; Shirayama et al., 2012; Woodhouse et al., 2018). Additionally, it is becoming clear that a large proportion of the autosomal DNA in *C. elegans* can be divided into active domains, containing H3K36me3 and germline expressed genes and regulated domains, containing H3K27me3 marked nucleosomes and silent genes (Evans et al., 2016; Liu et al., 2011). These domains are largely stable through development, including in the adult germline (Evans et al., 2016; Rechtsteiner et al., 2010). We examined the influence of these three types of chromatin on the response of TEs to mutation accumulation. TEs in regulated domains and in classical heterochromatin showed no significant overall increase in expression (Figure 5A, B). The rare examples of *N*.1 lines showing significantly higher expression of TEs in classical heterochromatin corresponded to lines in which Tc1 reactivation occurred, consistent with enrichment of Tc1 elements within these regions (McMurchy et al., 2017). Contrastingly, TEs in active domains had significantly higher expression in *N*.1 than *N*>1 lines (Wilcoxon unpaired test, *p* = 0.02). There was also a trend towards decreased median expression with increasing population size although this was on the border of significance (Jh, *p* = 0.07; LR, *p =* 0.08; Figure 5C).

**Figure 5.**
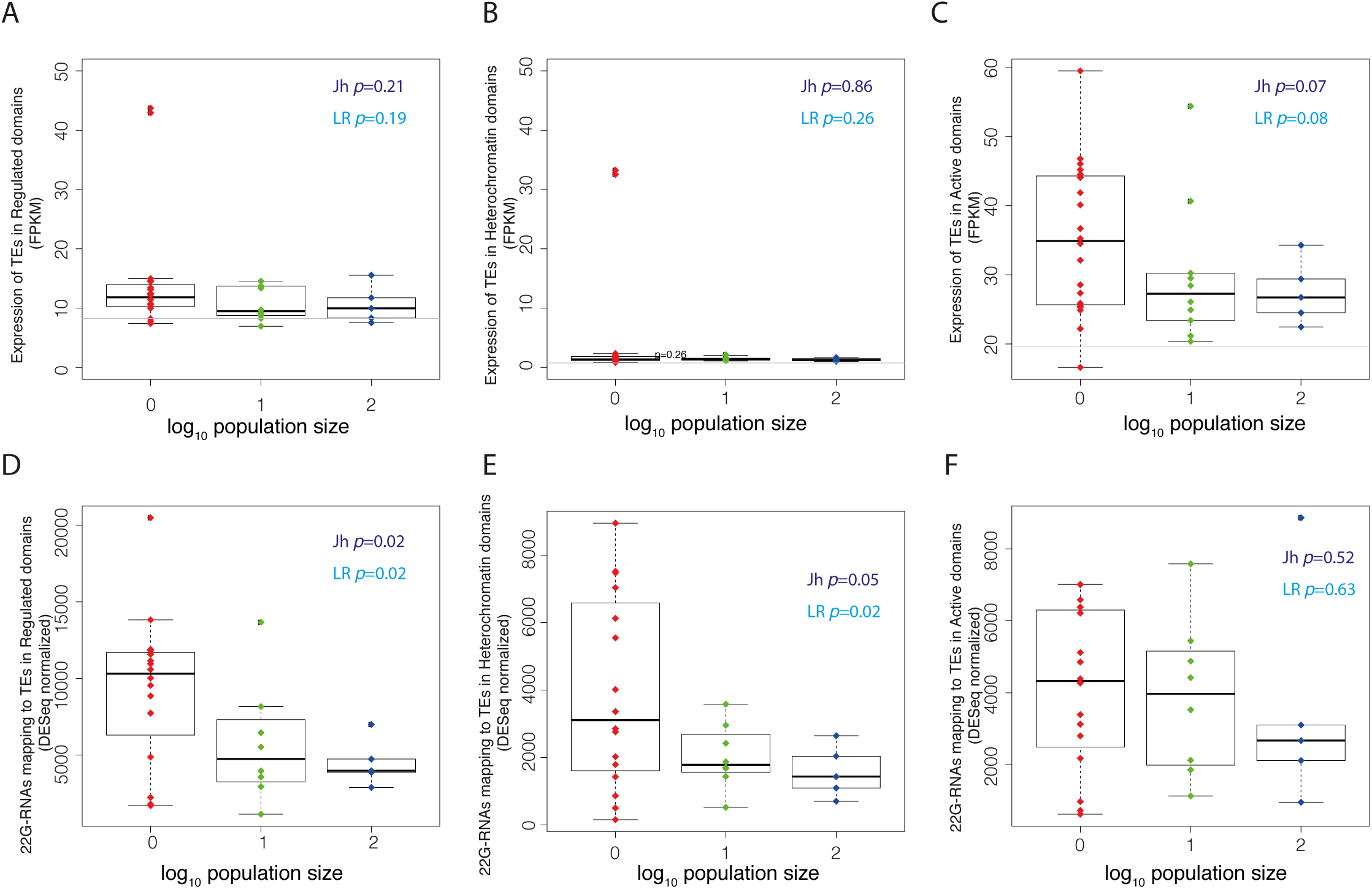
Chromatin environment controls alterations in piRNA mediated silencing in MA lines. **A, B, and C** Expression changes in TEs in Regulated (H3K27me3), Classic Heterochromatin (H3K9me2) and Active (H3K36me3) chromatin domains. *P*-values are Wilcoxon unpaired test. **D, E, and F** Changes in TE-mapping 22Gs in Regulated, Classic Heterochromatin and Active domains.

We next examined 22G-RNAs mapping to TEs across different chromatin domains. TEs in repressed domains showed significantly increased levels of 22G-RNAs in *N*.1 lines relative to *N*.10 and *N*.100. However, in active domains there was no significant change in 22G-RNA levels (Figure 5D-F). Thus, increased 22G-RNAs occur predominantly in TEs within repressed chromatin.

### AT-rich sequences in TEs reduce the generation of 22G-RNAs

A recent study has demonstrated that silencing of both transgenes and endogenous genes by 22G-RNAs is inhibited by a high content of periodic repeats of AT-rich sequences, known as PATCs (Frøkjær-Jensen et al., 2016). We tested how PATC density within TEs was associated with changes in their expression under reduced selection. High PATC density corresponded to reactivation of TEs (Jh, *p* = 0.02; LR, *p* = 0.02) whereas TEs with low PATC density did not show an increase in expression (Figure 6A). Contrastingly, only TEs with low PATC density showed significantly increased 22G-RNAs in *N*.1 lines relative to *N*.10 and *N*.100 (Jh, *p* = 0.02; LR, *p* = 0.01; Figure 6B). Importantly this effect was specific to PATC sequences as GC-content alone had no significant effect on either TE expression or small RNA generation (Supplemental Figure 4A, B). We conclude that low PATC density is required for 22G-RNA generation, which may be required to restrain TE activation. We tested whether the chromatin environment modulated the effect of PATC sequences on TE reactivation in MA lines. Importantly, PATC content was similar in TEs across active, classic heterochromatin and regulated domains (Supplemental Figure 4C). 22G-RNAs were significantly increased in low PATC regions within regulated domains (Jh, p = 0.02; LR, *p* = 0.11) and classical heterochromatin domains (Jh, *p* = 0.06; LR, *p* = 0.04) but not in active domains (Figure 6C, D). In contrast, there were no significant changes in 22G-RNA levels in high PATC regions within these domains (Figure 6C, D). Thus, the generation of increased 22G-RNAs against TEs in MA lines is associated with both low PATC content and a repressive chromatin environment.

**Figure 6.**
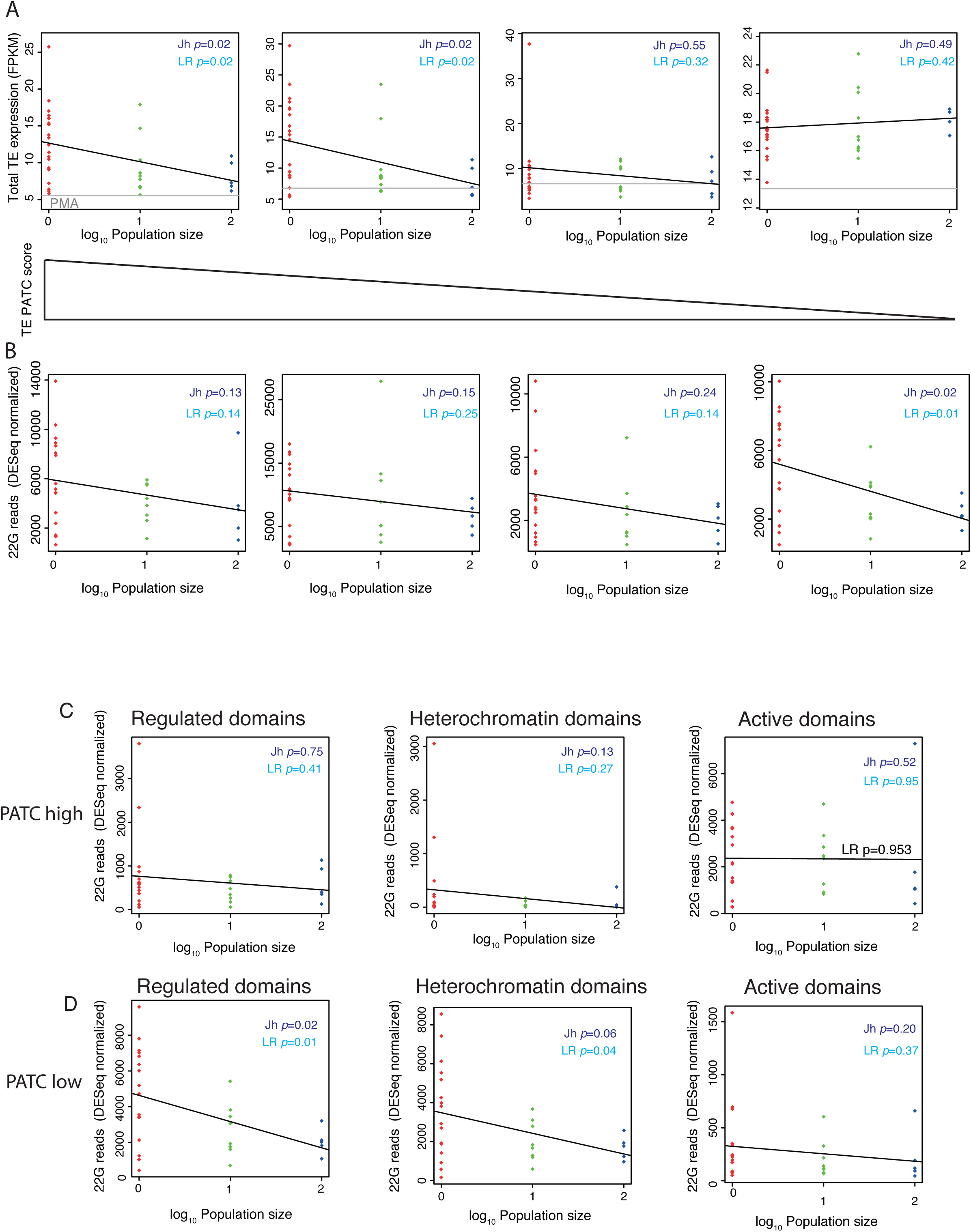
**PATC content influences 22G-RNA generation in MA lines A** Expression differences in TEs in four equal-size bins of 21 TEs with decreasing PATC content. **B** 22G-RNAs across the bins used in A. **C** and **D** Stratification of bins from A and B into regulated (H3K27me3) chromatin, active (H3K36me3) chromatin, and heterochromatin (H3K9me2) domains. High PATC is the top bin and low PATC is the lowest bin.

## Discussion

Our analysis of how the interplay between TE expression and TE silencing factors changes over 400 generations at small population sizes provides the first clear view of how TE expression diverges under reduced selection in animals. Additionally, closer analysis of how TE control mechanisms are affected in the MA lines offers new insight into the fundamental mechanisms of TE silencing in *C. elegans*, underlining the ability of experimental evolutionary studies to derive fundamental molecular insights. Here we discuss each of these aspects of our work in turn.

### The effect of selection on TE expression

Here, we demonstrate that in *C. elegans*, TEs drift to higher expression under conditions of strong genetic drift and reduced efficiency of selection. This is consistent with the action of purifying selection on TE expression. Only a single TE, Vingi-1, showed the opposite trend of lower expression in *N*.1 lines relative to the *N*.10 or *N*.100 lines (Figure 1B). However, the expression of this element was actually lower in the ancestral control than in the *N*.10 or *N*.100 lines. Thus, the significance of this observation is unclear. Our observations suggest that the expression of most TEs is largely detrimental and TE expression is under purifying selection. It remains a formal possibility that TE activation is beneficial under fluctuating environmental conditions or low fitness as opposed to the stable environment of the laboratory.

An important point raised by our results is that that not all expression increases of TEs are linked to increasing copy-number; indeed many lines with very high expression of specific TEs display no evidence of increased copy-number at all. This suggests that many TEs replicate inefficiently in *C. elegans* such that even very large increases in expression levels do not automatically result in increased copy-numbers. This also implies that TE expression may be detrimental without directly posing a threat to genome integrity, potentially through effects on endogenous gene expression networks or through toxicity of repetitive RNA within the cell (Simon et al., 2014).

Phenotypic analyses of previous *C. elegans* MA experiments suggest that the decline in fitness in *N* = 1 lines results primarily from a few mutations with large effects (Estes et al., 2004; Halligan et al., 2003; Katju et al., 2015, 2018; Keightley and Cabellero, 1997). Similar results have been obtained in experimental evolution studies in *D. melanogaster* (Ávila and García-Dorado, 2002) and bacteria (Dillon and Cooper, 2016; Heilbron et al., 2014). In our study, purifying selection at larger population sizes (*N*.10 and *N*.100) would eliminate such large-effect mutations and indeed, our *N*.10 and N. 100 lines exhibited no evidence of fitness reduction over the course of successive bottlenecking for 409 generations (Katju et al., 2015, 2018). Analysis of gene expression data from MA experiments from *C. elegans*, *D. melanogaster* and *S. cerevisiae* concluded that large effect mutations are also responsible for changes in protein-coding gene expression (Hodgins-Davis et al., 2015). However, our results for TE expression do not seem to fit with this model because overall TE expression, which is largely driven by Turmoil2 and non-LTR elements (see Figure 1), increases gradually with time across the *N*.1 lines and is also increased, although less so, in *N*.10 and *N*.100 lines. The expression of Tc1 is an exception to the overall trend as it is not affected in *N*.10 or *N*.100 lines but a small number of *N*.1 lines show markedly increased Tc1 expression. Thus, Tc1 reactivation may be dominated by a few mutations with large effect. This difference might be related to the different mechanism of silencing of Tc1 compared to Turmoil2 elements as discussed further below.

### Weakened piRNA silencing is responsible for increased expression of TEs under relaxed selection

Our investigations of the molecular mechanisms behind reduced silencing of TEs in MA lines strongly suggest that defective piRNA silencing is a major culprit. Only piRNA-targeted TEs show increased expression in MA lines, and, whilst piRNAs themselves do not seem to change significantly in MA lines, the levels of 22G-RNAs that act as effectors of piRNA silencing are perturbed at the Turmoil2 TEs that show increased expression. Why is piRNA silencing so vulnerable to mutation accumulation? TE silencing and TE activation in organisms are likely in a precarious equilibrium due to a constant evolutionary arms race between TEs and their host genome. As a result, many mutations could converge on the piRNA pathway to throw TE silencing out of balance.

### New insights into the role of chromatin in the piRNA pathway

Our analysis of how mutation accumulation affects TE silencing provides novel insights into how the chromatin environment of TEs might affect piRNA-mediated silencing in *C. elegans* (Figure 7). Importantly, TEs in repressive chromatin are much less prone to reactivation than those in active chromatin regions. Mechanistically this is because 22G-RNAs in repressive chromatin regions are stable or even increased, whilst 22G-RNAs mapping to TEs in active regions are reduced in MA lines with increased expression. This result may also explain why reactivation of Tc1 elements occurs less frequently than Turmoil2 elements, because Tc1 elements are predominantly located in repressed domains and are therefore silenced more robustly.

**Figure 7.**
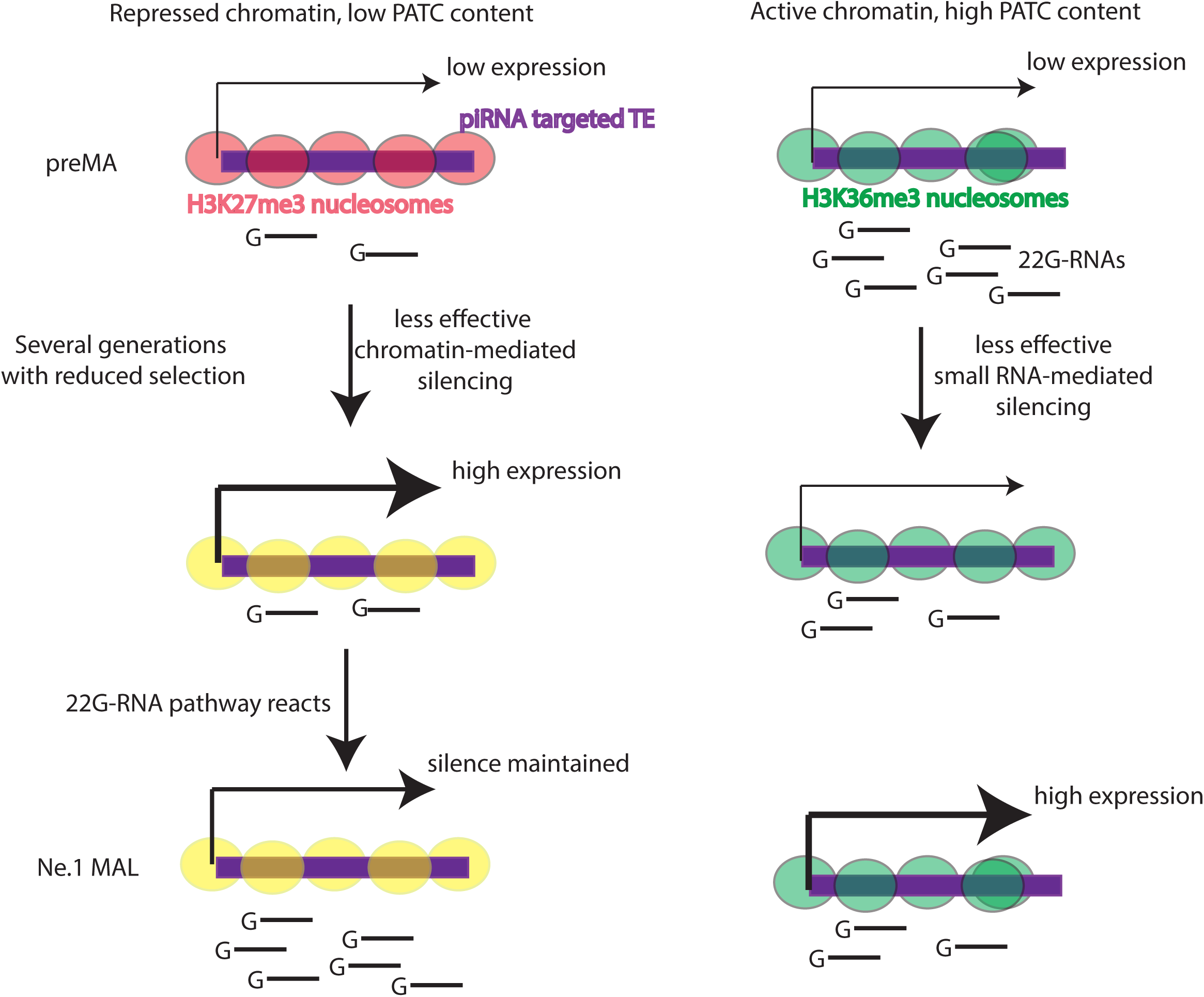
**Chromatin and PATC together influence TE control in MA lines** Model for how chromatin environment contributes to TE desilencing in MA lines. Left-repressed chromatin domains, taking H3K27me3 enriched domains as an example. Right, Active chromatin domains.

What is the mechanism whereby silencing memory is supported in repressed chromatin regions? The simplest possibility is that silencing and generation of 22G-RNAs are directly promoted by repressive chromatin modifications. In line with this possibility, a mutually reinforcing loop between H3K9 methylation and small RNAs is well documented in fission yeast (Bühler, 2009), and H3K9 methylation factors contribute to silencing of transgenes in *C. elegans* (Ashe et al., 2012; McMurchy et al., 2017; Shirayama et al., 2012; Woodhouse et al., 2018) although the situation is more complicated for endogenous genes (Ni et al., 2014). However, our observations hold equally well for H3K27me3-repressed chromatin, which has not been directly linked to 22G-RNA silencing. We propose therefore that the nuclear small RNA pathway responds differently depending on whether surrounding genes are active or repressed to detect and quell aberrant gene activation. This model will be of interest for further mechanistic investigation of small-RNA mediated silencing in *C. elegans*.

## Additional Files

R code and input files required to generate the plots in the figures will be released along with final publication and are available upon reasonable request to

## Supplemental Figure Legends

**Supplemental Figure 1.**
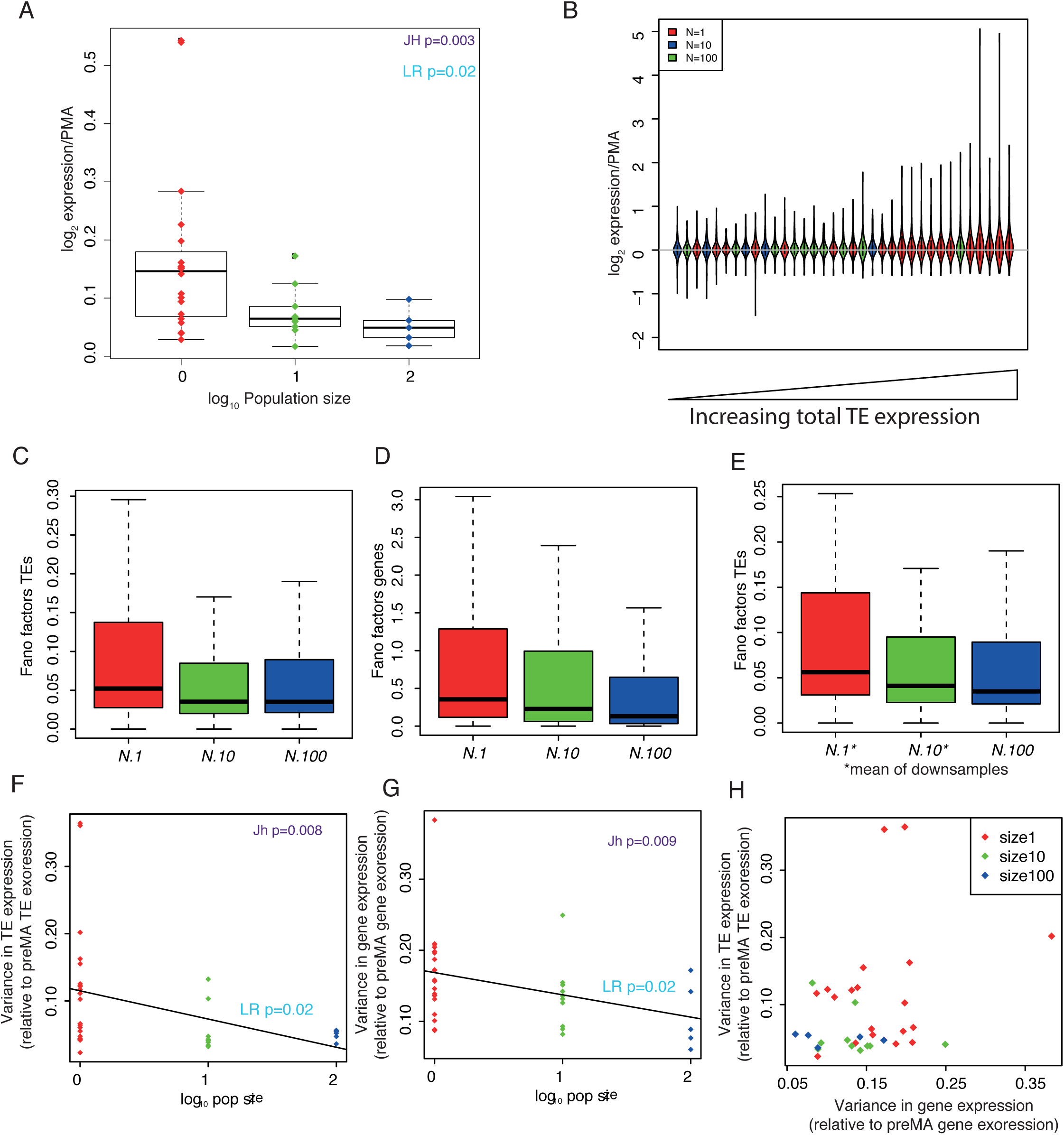
**A** Mean change in expression relative to the ancestral control across all TEs for each line. **B** Violin plots showing distribution of changes in TE expression across all TEs for each line. Lines are ordered by increasing mean TE expression change (L to R). **C** Fano factor of individual TE transcript levels across all lines of the indicated population size. **D** Fano factor in individual protein-coding gene transcript levels across all lines of the indicated population size. **E** Fano factor of TE transcript levels in 1,000 samples of five *N*.1 lines, 253 samples of five *N*.10 lines [the maximum] and all five *N*.100 lines. **F** Total variance in transcript level differences between each TE and its corresponding value in the starting population across lines of different population size. **G** Total variance of transcript level differences between each protein-coding gene and the starting population in lines of the indicated population size. Boxplots for A-D are as in Figure 1D. **H** Variance in TE transcript changes relative to starting population compared to the variance in protein-coding gene transcript changes in the same line.

**Supplemental Figure 2.**
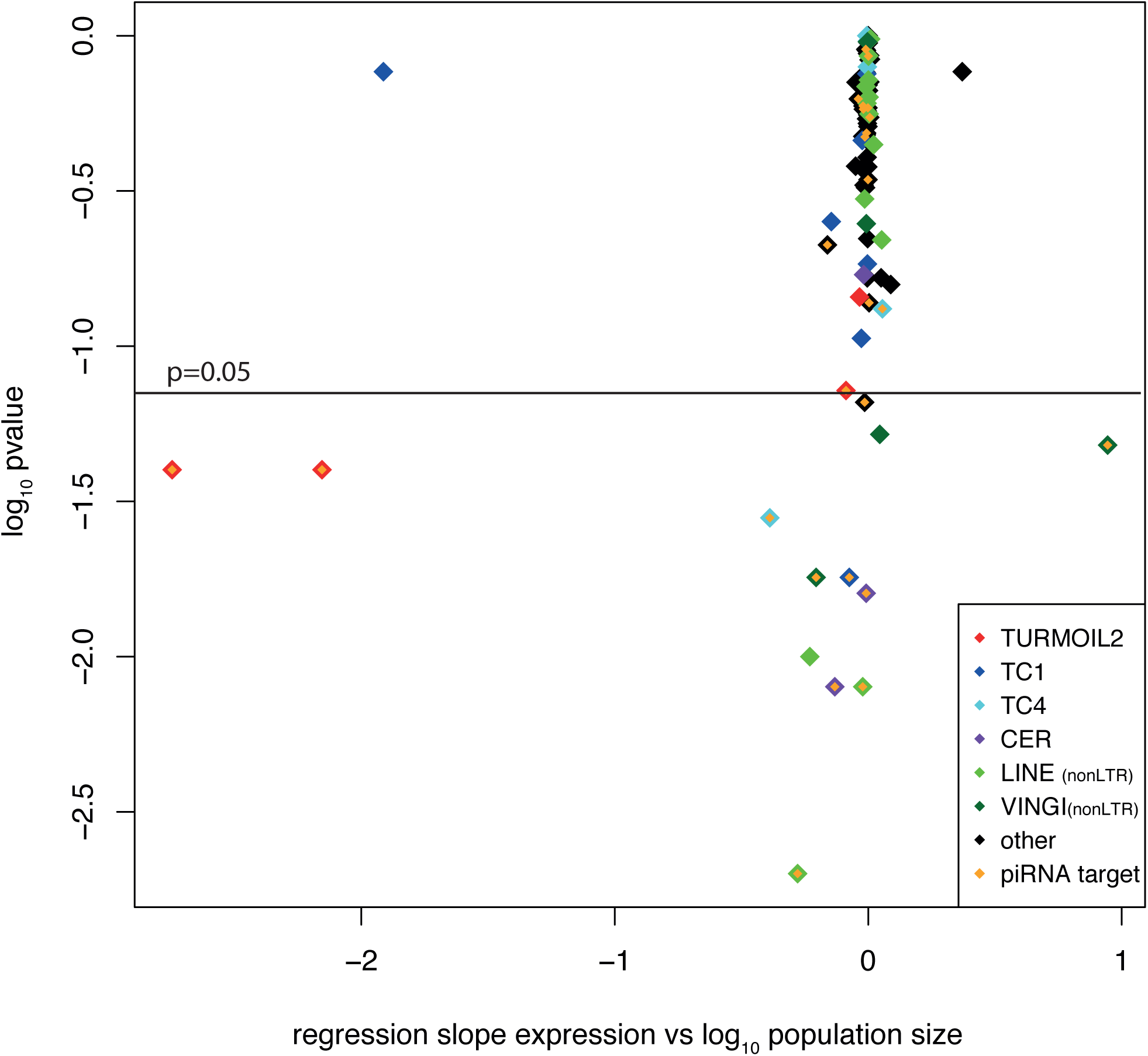
Volcano plot showing the logarithm (base 10) *p*-value of a linear model relating expression of TEs to population size on the y-axis to the gradient of the linear model on the x-axis.

**Supplemental Figure 3.**
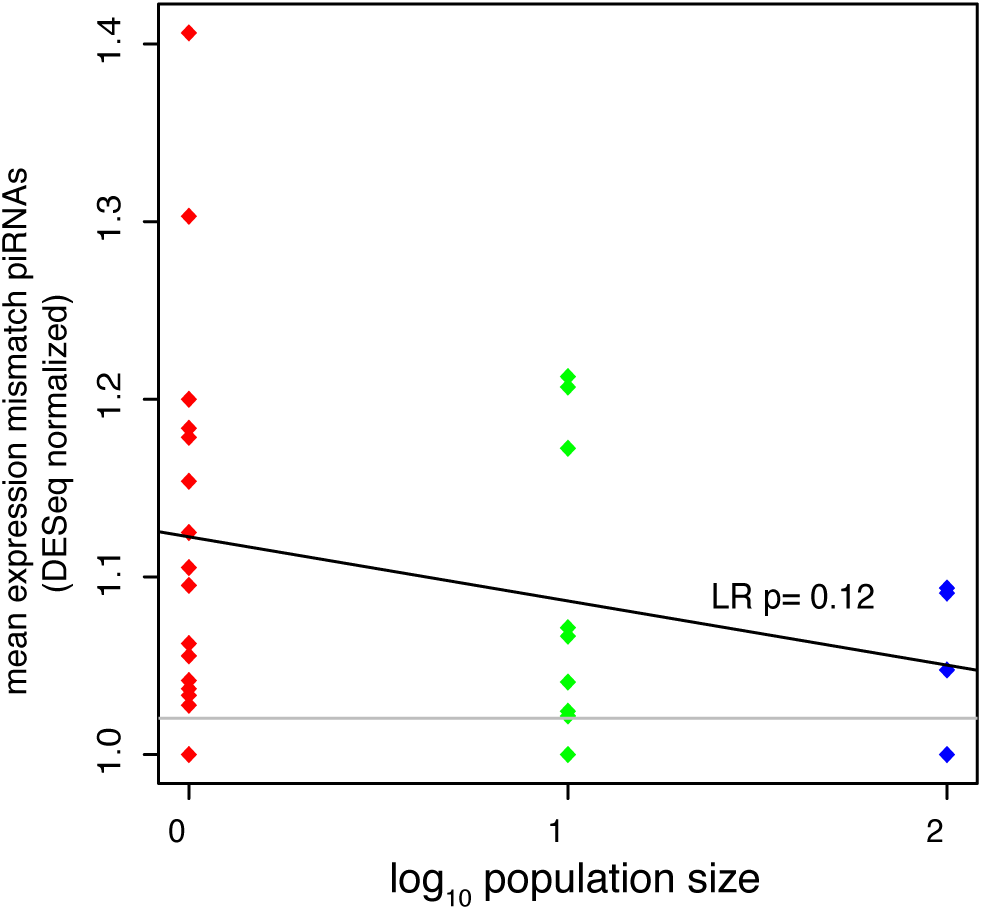
Total piRNA reads showing one base pair mismatch to the reference sequence divided by the total number of mismatched loci, in MA lines across different population sizes.

**Supplemental Figure 4.**
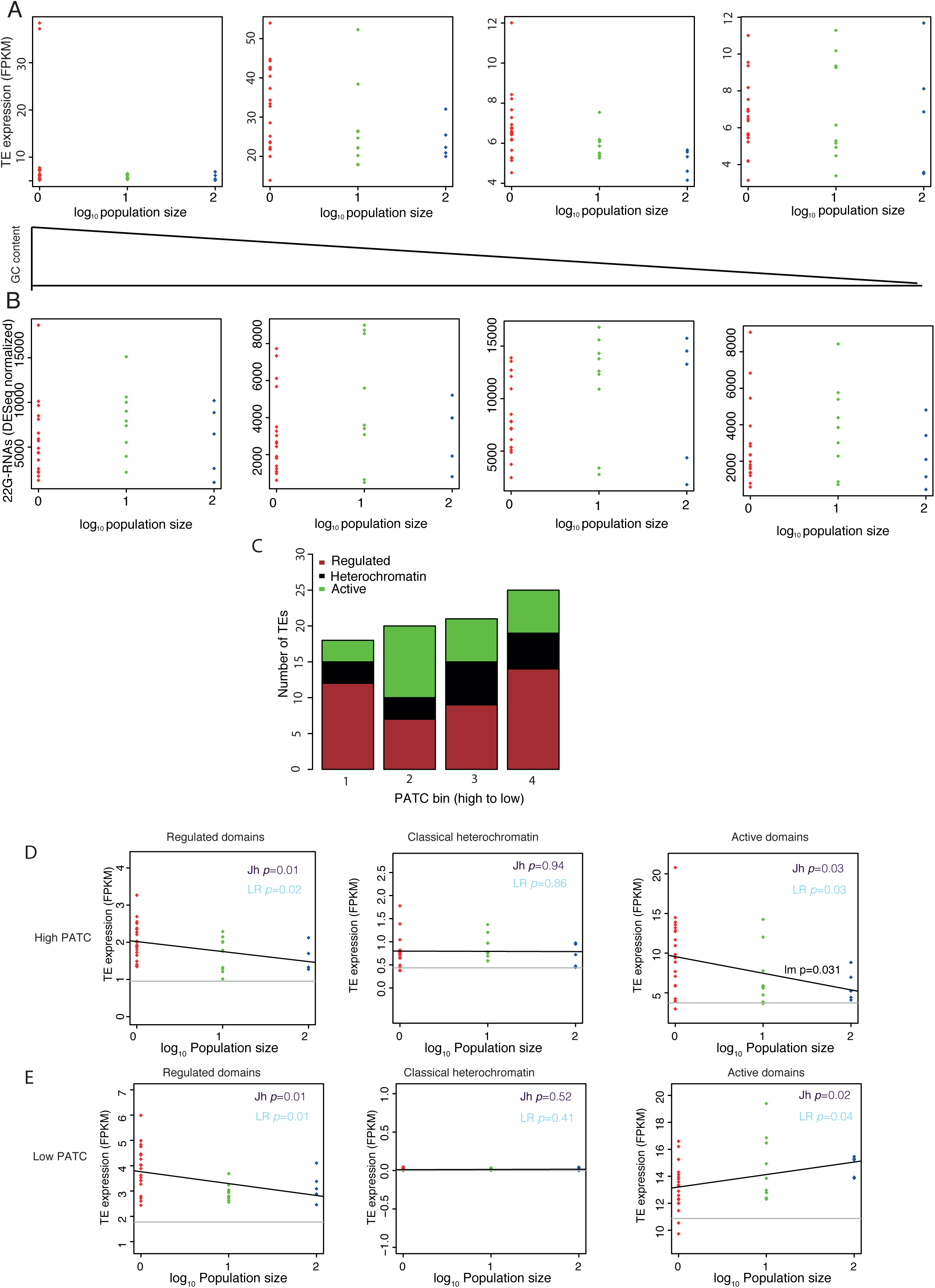
**A** GC content does not affect reactivation of TEs in the *N*.1 lines. Bins with high to low GC content, left to right. **B** GC content does not affect the change in 22G-RNA levels. Bins with high to low GC content, left to right. **C** No clear difference in the proportion of TEs from different chromatin domains within bins of different PATC content. **D** Expression of TEs from regulated, heterochromatic and active chromatin regions in the top PATC bin. **E** Expression of TEs from regulated, heterochromatic and active chromatin regions in the lowest PATC bin.

## Methods

### Spontaneous Mutation Accumulation (MA) Experiment at Varying Population Sizes and Its Theoretical Underpinnings

The descendants of a single wild-type Bristol (N2) hermaphrodite were used to establish 35 MA lines. Twenty MA lines were propagated through single individual descent, while ten lines were bottlenecked to ten individuals, and five lines were bottlenecked to 100 individuals every generation (Katju et al., 2015, 2018; Konrad et al., 2018)(Katju et al., 2015, 2018; Konrad et al. 2018). The size of these bottlenecks was ensured through careful counting of randomly chosen L4 larvae selected to establish a new generation every four days. All populations were grown on NGM plates (Nematode Growth Medium) at 20°C (Figure 1A). The lines were propagated through 409 generations or until extinction under standard laboratory conditions. Populations of size *N* = 1 (*N*.1 lines) and 10 (*N*.10 lines) were maintained on 60×15 mm petri dishes seeded with 250*µl* of *Escherichia coli* (OP50) in YT medium. *N* = 100 (or *N*.100) populations were housed on 90×15 mm petri dishes seeded with 750*µl* OP50 in YT medium.

The fitness effects of mutations range continuously from lethal to deleterious to neutral to beneficial. In small populations, beneficial mutations can be lost and detrimental mutations fixed by random chance events (genetic drift). The loss or fixation of mutations depend upon both their selection coefficients (*s*) and the effective population size, *N_e_*. It has been shown that for sexually reproducing diploids, the dynamics of mutations with |s| << 1/2*N_e_* are dominated by random genetic drift(Kimura, 1962). Therefore, small populations subjected to attenuated selection and an increased magnitude of genetic drift can potentially accumulate mutations with extremely large effects in addition to ones with moderate to very slight effects. With increasing population size, the efficiency of natural selection is increased. The differences in populations size in this MA experiment alters the relative importance of genetic drift versus natural selection in the fixation or loss of mutations, with genetic drift having the greatest influence in *N* = 1 lines and natural selection having greater influence in populations that were bottlenecked at 10 and 100 individuals each generation.

### RNA Library Preparation, Sequencing, and Analysis of Transcript Abundance

The library preparation and RNA-sequencing procedures have previously been described in detail (Konrad et al., 2018). Briefly, we isolated one, two, and three individuals each of the *N*.1, *N*.10, and *N*.100 lines, respectively. These 55 worms, as well as one individual from the ancestral population, were each sequestered to NGM plates seeded with OP50, where they were allowed to self-fertilize and reproduce at 20°C. Three offspring worms at the L4 larval stage were isolated from each of the F_1_ populations to serve as biological replicates. These 168 individual worm samples were allowed to reproduce for three generations to yield enough tissue for RNA extraction. A standard bleaching protocol was used to collect gravid eggs from adults in order to generate synchronized populations of L1 larvae. Total RNA was isolated from L1 larvae via the Qiagen RNeasy Mini Kit. The Nanodrop 2000, Qubit 3.0 Fluorometer, and an Agilent RNA Analyzer were used to evaluate the quality of the RNA samples, and an Illumina TruSeq RNA library Prep Kit v2 was used with standard procedures to prepare the RNA sequencing libraries for each sample at the Texas A&M University Genomics and Bioinformatics Services Center. The RNA was fragmented and Illumina adapters were annealed for amplification. Size selected cDNA fragments were isolated via a Qiagen Gel Extraction Kit. Finally, sequencing was performed on the Illumina HiSeq 4000 platform with default quality filters.

Demultiplexing and prefiltering of the sequencing reads was performed based on default Illumina QC protocols. Reads containing abnormally short insert lengths were removed, and adapters were discarded from the reads. The raw RNA-sequencing reads in fastq format were aligned to the protein-coding transcriptome of *C. elegans* (Wormbase reference N2 genome version WS247) using TopHat(Trapnell et al., 2009) via the “very sensitive” bowtie2 algorithm with a maximum of one mismatch in the anchor region for each spliced alignment and a minimum and maximum intron length of 20 and 3,000 bp, respectively. Cufflinks(Trapnell et al., 2010) with default settings and gene annotations from the N2 genome version WS247 was used to estimate the relative transcript abundance for each protein-coding gene. All following analyses were focused on FPKM values calculated on the per gene level. The relative transcript abundances (FPKM) from the three biological replicates for each original sample were averaged to get mean relative transcript abundance for each gene in that sample.

### Small RNA sequencing

MA lines were synchronised using hypochlorite treatment and embryos were isolated after 12 hours. RNA was extracted using trizol and small RNA libraries were prepared using the Illumina Small RNA sequencing kit as described previously(Sarkies et al., 2015). Small RNAs were aligned to a genome built using bowtie from a fasta file containing all piRNAs, miRNAs, ncRNAs and genes including TEs, extracted from Wormbase (WS264; ce11), requiring perfect mapping. Reads mapping to the sense strand of ncRNAs and miRNAs were extracted using bedtools intersect –c –S. We used DEseq using ncRNAs and miRNAs to extract size factors. 22G-RNAs mapping to TEs and genes were extracted using a custom Perl script and the number of 22G-RNAs mapping antisense to each gene and TE was then counted using bedtools intersect –c-s. The 22G-RNAs were then normalized to the size factors from ncRNAs and miRNAs combined.

### TE copy-number analysis

DNA sequence reads were aligned to a genome built using bowtie2-build from TE consensus sequences extracted from repbase combined with all coding sequences. Bowtie2 was used to map PE reads to this genome and the read count mapping to each CDS or TE was obtained using bedtools intersect –c.

### Computational analysis of TE expression and small RNA analysis

TEs from WS264 were annotated using Repeatmasker(Smit et al., 2017). All data analysis was conducted using the R environment for statistical analysis (www.Rproject.com). Details of the individual analyses are documented in the R markdown file accompanying this manuscript (additional file 1), with the raw data tables required to run these programs in as a zipped file (additional file 2). Previously published datasets containing Chromatin domain annotations from Early Embryo ChiP-Seq were taken from Evans et al. (2016) updated to WS264 using liftover (https://genome.ucsc.edu/). Small RNA sequencing data from reactivation of small RNA pathways in the presence or absence of piRNAs was taken from Phillips et al. (2015) and aligned to the *C. elegans* genome as described above. The average PATC score for each TE was calculated by taking the average PATC score across the element from per-base sliding window genome-wide PATC scores from (Frøkjær-Jensen et al., 2016).

## REFERENCES

Adrion, J.R., Song, M.J., Schrider, D.R., Hahn, M.W., and Schaack, S. (2017). Genome-wide estimates of transposable element insertion and deletion rates in *Drosophila melanogaster*. Genome Biol. Evol. 9, 1329–1340.

de Albuquerque, B.F.M., Placentino, M., and Ketting, R.F. (2015). Maternal piRNAs are essential for germline development following de novo establishment of endo-siRNAs in *Caenorhabditis elegans*. Dev. Cell 34, 448–456.

Ashe, A., Sapetschnig, A., Weick, E.M., Mitchell, J., Bagijn, M.P., Cording, A.C., Doebley, A.L., Goldstein, L.D., Lehrbach, N.J., Le Pen, J., et al. (2012). PiRNAs can trigger a multigenerational epigenetic memory in the germline of *C. elegans*. Cell 150, 88–99.

Ávila, V., and García-Dorado, A. (2002). The effects of spontaneous mutation on competitive fitness in *Drosophila melanogaster*. J. Evol. Biol. 15, 561–566.

Bagijn, M.P., Goldstein, L.D., Sapetschnig, A., Weick, E.M., Bouasker, S., Lehrbach, N.J., Simard, M.J., and Miska, E.A. (2012). Function, targets, and evolution of *Caenorhabditis elegans* piRNAs. Science (80-.). 337, 574–578.

Bast, J., Jaron, K.S., Schuseil, D., Roze, D., and Schwander, T. (2019). Asexual reproduction reduces transposable element load in experimental yeast populations. Elife 8.

Batista, P.J., Ruby, J.G., Claycomb, J.M., Chiang, R., Fahlgren, N., Kasschau, K.D., Chaves, D.A., Gu, W., Vasale, J.J., Duan, S., et al. (2008). PRG-1 and 21U-RNAs interact to form the piRNA complex required for fertility in *C. elegans*. Mol. Cell 31, 67–78.

Bégin, M., and Schoen, D.J. (2006). Low impact of germline transposition on the rate of mildly deleterious mutation in *Caenorhabditis elegans*. Genetics 174, 2129–2136.

Bégin, M., and Schoen, D.J. (2007). Transposable elements, mutational correlations, and population divergence in *Caenorhabditis elegans*. Evolution (N. Y). 61, 1062–1070.

Beltran, T., Barroso, C., Birkle, T.Y., Stevens, L., Schwartz, H.T., Sternberg, P.W., Fradin, H., Gunsalus, K., Piano, F., Sharma, G., et al. (2019). Comparative epigenomics reveals that rna polymerase ii pausing and chromatin domain organization control nematode piRNA biogenesis. Dev. Cell 48, 793–810.e6.

Billi, A.C., Freeberg, M.A., Day, A.M., Chun, S.Y., Khivansara, V., and Kim, J.K. (2013). a conserved upstream motif orchestrates autonomous, germline-enriched expression of *Caenorhabditis elegans* piRNAs. PLoS Genet. 9.

Brennecke, J., Aravin, A.A., Stark, A., Dus, M., Kellis, M., Sachidanandam, R., and Hannon, G.J. (2007). Discrete small rna-generating loci as master regulators of transposon activity in *Drosophila*. Cell 128, 1089–1103.

Buckley, B.A., Burkhart, K.B., Gu, S.G., Spracklin, G., Kershner, A., Fritz, H., Kimble, J., Fire, A., and Kennedy, S. (2012). A nuclear Argonaute promotes multigenerational epigenetic inheritance and germline immortality. Nature 489, 447–451.

Bühler, M. (2009). RNA turnover and chromatin-dependent gene silencing. Chromosoma 118, 141–151.

Cecere, G., Zheng, G.X.Y., Mansisidor, A.R., Klymko, K.E., and Grishok, A. (2012). Promoters recognized by forkhead proteins exist for individual 21U-RNAs. Mol. Cell 47, 734–745.

Chuong, E.B., Elde, N.C., and Feschotte, C. (2017). Regulatory activities of transposable elements: From conflicts to benefits. Nat. Rev. Genet. 18, 71–86.

Das, P.P., Bagijn, M.P., Goldstein, L.D., Woolford, J.R., Lehrbach, N.J., Sapetschnig, A., Buhecha, H.R., Gilchrist, M.J., Howe, K.L., Stark, R., et al. (2008). Piwi and piRNAs act upstream of an endogenous siRNA pathway to suppress tc3 transposon mobility in the *Caenorhabditis elegans* germline. Mol. Cell 31, 79–90.

Denver, D.R., Morris, K., Streelman, J.T., Kim, S.K., Lynch, M., and Thomas, W.K. (2005). The transcriptional consequences of mutation and natural selection in Caenorhabditis elegans. Nat. Genet. 37, 544–548.

Dillon, M.M., and Cooper, V.S. (2016). The fitness effects of spontaneous mutations nearly unseen by selection in a bacterium with multiple chromosomes. Genetics 204, 1225–1238.

Estes, S., Phillips, P.C., Denver, D.R., Thomas, W.K., and Lynch, M. (2004). Mutation accumulation in populations of varying size: the distribution of mutational effects for fitness correlates in *Caenorhabditis elegans*. Genetics 166, 1269–1279.

Evans, K.J., Huang, N., Stempor, P., Chesney, M.A., Down, T.A., and Ahringer, J. (2016). Stable *Caenorhabditis elegans* chromatin domains separate broadly expressed and developmentally regulated genes. Proc. Natl. Acad. Sci. 113, E7020--E7029.

Fontenla, S., Rinaldi, G., Smircich, P., and Tort, J.F. (2017). Conservation and diversification of small RNA pathways within flatworms. BMC Evol. Biol.

Frøkjær-Jensen, C., Jain, N., Hansen, L., Davis, M.W., Li, Y., Zhao, D., Rebora, K., Millet, J.R.M., Liu, X., Kim, S.K., et al. (2016). An abundant class of non-coding DNA can prevent stochastic gene silencing in the *C. elegans* germline. Cell 166, 343– 357.

Gu, W., Lee, H.-C., Chaves, D., Youngman, E.M., Pazour, G.J., Conte, D., and Mello, C.C. (2012). CapSeq and CIP-TAP identify Pol II start sites and reveal capped small RNAs as C. elegans piRNA precursors. Cell 151, 1488–1500.

Halligan, D.L., and Keightley, P.D. (2009). Spontaneous mutation accumulation studies in evolutionary genetics. Annu. Rev. Ecol. Evol. Syst. 40, 151–172.

Halligan, D., Peters, A., and Keightley, P. (2003). Estimating numbers of EMS-induced mutations affecting life history traits in *Caenorhabditis elegans* in crosses between inbred sublines. Genet. Res. 82, 191–205.

Heilbron, K., Toll-Riera, M., Kojadinovic, M., and MacLean, R.C. (2014). Fitness is strongly influenced by rare mutations of large effect in a microbial mutation accumulation experiment. Genetics 197, 981–990.

Hodgins-Davis, A., Rice, D.P., and Townsend, J.P. (2015). Gene expression evolves under a house-of-cards model of stabilizing selection. Mol. Biol. Evol. 32, 2130– 2140.

Imbeault, M., Helleboid, P.-Y., and Trono, D. (2017). KRAB zinc-finger proteins contribute to the evolution of gene regulatory networks. Nature 543, 550–554.

Katju, V., and Bergthorsson, U. (2019). Old trade, new tricks: insights into the spontaneous mutation process from the partnering of classical mutation accumulation experiments with high-throughput genomic approaches. Genome Biol. Evol. 11, 136–165.

Katju, V., Packard, L.B., Bu, L., Keightley, P.D., and Bergthorsson, U. (2015). Fitness decline in spontaneous mutation accumulation lines of *Caenorhabditis elegans* with varying effective population sizes. Evolution (N. Y). 69, 104–116.

Katju, V., Packard, L.B., and Keightley, P.D. (2018). Fitness decline under osmotic stress in *Caenorhabditis elegans* populations subjected to spontaneous mutation accumulation at varying population sizes. Evolution (N. Y). 72, 1000–1008.

Keightley, P.D., and Cabellero, A. (1997). Genomic mutation rates for lifetime reproductive output and lifespan in Caenorhabditis elegans. Proc. Natl. Acad. Sci. USA 94, 3823–3827.

Kimura, M. (1962). On the probability of fixation of mutant genes in a population. Genetics 47, 713–719

Konrad, A., Flibotte, S., Taylor, J., Waterston, R.H., Moerman, D.G., Bergthorsson, U., and Katju, V. (2018). Mutational and transcriptional landscape of spontaneous gene duplications and deletions in *Caenorhabditis elegans*. Proc. Natl. Acad. Sci. 115, 7386–7391.

Landry, C.R., Lemos, B., Rifkin, S.A., Dickinson, W.J., and Hartl, D.L. (2007). Genetic properties influencing the evolvability of gene expression. Science (80-.). 317, 118–121.

Liu, T., Rechtsteiner, A., Egelhofer, T.A., Vielle, A., Latorre, I., Cheung, M.-S., Ercan, S., Ikegami, K., Jensen, M., Kolasinska-Zwierz, P., et al. (2011). Broad chromosomal domains of histone modification patterns in *C. elegans*. Genome Res. 21, 227–236.

Luteijn, M.J., van Bergeijk, P., Kaaij, L.J.T., Almeida, M.V., Roovers, E.F., Berezikov, E., and Ketting, R.F. (2012). Extremely stable Piwi-induced gene silencing in *Caenorhabditis elegans*. EMBO J. 31, 3422–3430.

McMurchy, A.N., Stempor, P., Gaarenstroom, T., Wysolmerski, B., Dong, Y., Aussianikava, D., Appert, A., Huang, N., Kolasinska-Zwierz, P., Sapetschnig, A., et al. (2017). A team of heterochromatin factors collaborates with small RNA pathways to combat repetitive elements and germline stress. Elife 6.

Mondal, M., Klimov, P., and Flynt, A.S. (2018). Rewired RNAi-mediated genome surveillance in house dust mites. PLoS Genet. 14.

Ni, J.Z., Chen, E., and Gu, S.G. (2014). Complex coding of endogenous siRNA, transcriptional silencing and H3K9 methylation on native targets of germline nuclear RNAi in *C. elegans*. BMC Genomics 15, 1157.

Pak, J., and Fire, A. (2007). Distinct populations of primary and secondary effectors during RNAi in C. *elegans*. Science (80-.). 315, 241–244.

Percharde, M., Lin, C.-J., Yin, Y., Guan, J., Peixoto, G.A., Bulut-Karslioglu, A., Biechele, S., Huang, B., Shen, X., and Ramalho-Santos, M. (2018). A LINE1-nucleolin partnership regulates early development and ESC Identity. Cell 174, 391–405.e19.

Phillips, C.M., Brown, K.C., Montgomery, B.E., Ruvkun, G., and Montgomery, T.A. (2015). PiRNAs and piRNA-dependent siRNAs protect conserved and essential *C. elegans* genes from misrouting into the RNAi Pathway. Dev. Cell 34, 457–465.

Rebollo, R., Romanish, M.T., and Mager, D.L. (2011). Transposable Elements: an abundant and natural source of regulatory sequences for host genes. Annu. Rev. Genet. 46, 120913153128008.

Rechtsteiner, A., Ercan, S., Takasaki, T., Phippen, T.M., Egelhofer, T.A., Wang, W., Kimura, H., Lieb, J.D., and Strome, S. (2010). The histone H3K36 methyltransferase MES-4 acts epigenetically to transmit the memory of germline gene expression to progeny. PLOS Genet. 6, e1001091.

Rifkin, S.A., Houle, D., Kim, J., and White, K.P. (2005). A mutation accumulation assay reveals a broad capacity for rapid evolution of gene expression. Nature 438, 220–223.

Ruby, J.G., Jan, C., Player, C., Axtell, M.J., Lee, W., Nusbaum, C., Ge, H., and Bartel, D.P. (2006). Large-scale sequencing reveals 21U-RNAs and additional microRNAs and endogenous siRNAs in *C. elegans*. Cell 127, 1193–1207.

Sarkies, P., Selkirk, M.E., Jones, J.T., Blok, V., Boothby, T., Goldstein, B., Hanelt, B., Ardila-Garcia, A., Fast, N.M., Schiffer, P.M., et al. (2015). Ancient and novel small RNA pathways compensate for the loss of piRNAs in Multiple Independent Nematode Lineages. PLoS Biol. 13.

Shen, E.-Z., Chen, H., Ozturk, A.R., Tu, S., Shirayama, M., Tang, W., Ding, Y.-H., Dai, S.-Y., Weng, Z., and Mello, C.C. (2018). Identification of piRNA binding sites reveals the argonaute regulatory landscape of the *C. elegans* germline. Cell 172, 937–951.e18.

Shirayama, M., Seth, M., Lee, H.-C., Gu, W., Ishidate, T., Conte, D., and Mello, C.C. (2012). piRNAs initiate an epigenetic memory of nonself RNA in the *C. elegans* germline. Cell 150, 65–77.

Simon, M., Sarkies, P., Ikegami, K., Doebley, A.L., Goldstein, L.D., Mitchell, J., Sakaguchi, A., Miska, E.A., and Ahmed, S. (2014). Reduced Insulin/IGF-1 signaling restores germ cell immortality to *Caenorhabditis elegans* Piwi mutants. Cell Rep. 7, 762–773.

Simonti, C.N., Pavličev, M., and Capra, J.A. (2017). Transposable Element exaptation into regulatory regions is rare, influenced by evolutionary age, and subject to pleiotropic constraints. Mol. Biol. Evol. 34, 2856–2869.

Siomi, M.C., Sato, K., Pezic, D., and Aravin, A.A. (2011). PIWI-interacting small RNAs: the vanguard of genome defence. Nat. Rev. Mol. Cell Biol. 12, 246.

Skinner, D.E., Rinaldi, G., Koziol, U., Brehm, K., and Brindley, P.J. (2014). How might flukes and tapeworms maintain genome integrity without a canonical piRNA pathway? Trends Parasitol.

Smit, A., Hubley, R., and Green, P. (2017). RepeatMasker Open-4.0.6 2013-2015. Http://Www.Repeatmasker.Org.

Szitenberg, A., Cha, S., Opperman, C.H., Bird, D.M., Blaxter, M.L., and Lunt, D.H. (2016). Genetic drift, not life history or RNAi, determine long-term evolution of Transposable Elements. Genome Biol. Evol. 8, 2964–2978.

Trapnell, C., Pachter, L., and Salzberg, S.L. (2009). TopHat: discovering splice junctions with RNA-Seq. Bioinformatics 25, 1105–1111.

Trapnell, C., Williams, B. a, Pertea, G., Mortazavi, A., Kwan, G., van Baren, M.J., Salzberg, S.L., Wold, B.J., and Pachter, L. (2010). How Cufflinks works. Nat. Biotechnol. 28, 511–515.

Wang, G., and Reinke, V. (2008). A *C. elegans* Piwi, PRG-1, regulates 21U-RNAs during spermatogenesis. Curr. Biol. 18, 861–867.

Weick, E., and Miska, E. a (2014). piRNAs : from biogenesis to function. Genes Dev. 1–41.

Woodhouse, R.M., Buchmann, G., Hoe, M., Harney, D.J., Low, J.K.K., Larance, M., Boag, P.R., and Ashe, A. (2018). Chromatin modifiers SET-25 and SET-32 Are required for establishment but not long-term maintenance of transgenerational epigenetic inheritance. Cell Rep. 25, 2259–2272.e5.

Yigit, E., Batista, P.J., Bei, Y., Pang, K.M., Chen, C.-C.G., Tolia, N.H., Joshua-Tor, L., Mitani, S., Simard, M.J., and Mello, C.C. (2006). Analysis of the *C. elegans* Argonaute family reveals that distinct Argonautes act sequentially during RNAi. Cell 127, 747–757.

